# Automated bioprocess feedback operation in a high throughput facility via the integration of a mobile robotic lab assistant

**DOI:** 10.1101/2022.01.13.476044

**Authors:** Lucas Kaspersetz, Saskia Waldburger, M.-Therese Schermeyer, Sebastian L. Riedel, Sebastian Groß, Peter Neubauer, M.-Nicolas Cruz-Bournazou

## Abstract

Biotechnological processes development is challenging due to the sheer variety of process parameters. For efficient upstream development parallel cultivation systems have proven to reduce costs and associated timelines successfully, while offering excellent process control. However, the degree of automation of such small scale systems is comparably low and necessary sample analysis requires manual steps. Although the subsequent analysis can be performed in a high-throughput manner, the integration of analytic devices remains challenging. Especially, when cultivation and analysis laboratories are spatially separated. Mobile robots offer a potential solution, but the implementation in research laboratories is not widely adopted. Our approach demonstrates the integration of a small scale cultivation system into a liquid handling station for an automated sample procedure. The samples are transferred via a mobile robotic lab assistant and subsequently analysed by a high-throughput analyzer. The process data is stored in a centralized database. The mobile robotic workflow guarantees a flexible solution for device integration and facilitates automation. Restrictions regarding spatial separation of devices are circumvented, enabling a modular platform throughout different laboratories. The presented cultivation platform is evaluated based on industrial relevant *E. coli* BW25113 high cell density fed-batch cultivation. Here its suitability for accelerating bioprocess development is proven. The necessary magnesium addition for reaching high cell densities in mineral salt medium is automated via a feedback operation loop. The feedback operation loop demonstrates the possibility for advanced control options. This study sets the foundation for a fully integrated facility with different cultivation scales sharing the same data infrastructure, where the mobile robotic lab assistant physically connects the devices.

## Introduction

he development of new biotechnological products is a laborious multi-stage process. The set-up of a final large-scale process generally takes about 5-10 years (1). The development cycle usually follows a sequential workflow, where throughput and information content follow opposite directions (2, 3). In order to shorten these development cycles and reduce costs, high-throughput (HT) robotic facilities have become inevitable for early stage upstream bioprocess development. Miniaturized and parallelized experiments, operated automatically in liquid handling stations (LHS) ensure the necessary throughput and reduce experimental effort (4–7). The smallest entity handled are microwell plates (MWP) with cultivation volumes in the range of 50 - 1000 μL or below. The main disadvantages of screening in MWP format is that they do not resemble industrial conditions, especially when operated under batch conditions. Although this issue has been solved and fed-batch operation is possible (8), limitations regarding scalability and process insights are still challenging (2). Therefore, many miniature bioreactor concepts, geometrically similar to large scale, have been developed and investigated throughout the last years. For a more detailed review see (2) or (3). The systems presented in the aforementioned publications represent industrially relevant conditions better than simple MWPs, but suffer from several drawbacks including low culture volumes, leading to small sample volumes, insufficient process flexibility to investigate all relevant parameters and limitations in individual process control. Furthermore, major drawbacks are very limited feeding strategies as well as the inability to reach industrially relevant high cell densities (HCD) (9). In a fed-batch process, the continuous feed of a growth limiting substrate avoids the formation of acetate due to overflow metabolism (10–12). Therefore, high cell densities with concentrations of ≥ 50 g L^−1^ can be routinely obtained for recombinant or non-recombinant *Escherichia coli* strains (13–15). Only a few studies have successfully challenged automated mini bioreactor systems for such high cell densities. (16) reported cell dry weights (CDW) up to ≥ 70 gL^−1^ in 10 mL disposable reactors. In a similar reactor type optical densities ≥ 100 have been reached (17). Nevertheless, constraints regarding low culture volume with adequate sample volumes, individual process control and feeding strategies with continuous feeding remain an issue. In contrast to high-throughput mini bioreactor systems, these constraints are typically negligible for small scale or bench scale cultivation systems, still being the standard in many research facilities. However, a decrease in throughput, manual sampling procedures and no subsequent sample processing are commonly the trade-off. Consequently, device integration concepts and automation tools should certainly be generalized, modularized and transferred to small scale and laboratory scale cultivation systems (18). Independent of their scale, cultivation systems have to be operated and controlled under optimal conditions. These automation tools must not be limited to the coupling of devices, but also necessary interfaces for data driven or model-based control strategies are required. Model-based tools have proven to be of utmost importance with regard to optimal experimental design (19) and reducing experimental effort, when limited prior knowledge about the strain or process is present (20, 21). For a recent review on model-based tools and their applications in bioprocess see (22) or (23). In order to make use of the aforementioned frameworks for optimal process control and deriving process specific parameters such as yields and growth rates, the acquisition of online, at-line and off-line data is mandatory. Especially in high-throughput scenarios, sample frequency to infere the phenotype of microbial cultivations can be very challenging (7). While a variety of on-line, in-line and at-line probes for real-time process analytics exist and essential substrates or metabolites can be estimated *in situ* (24), these methods such as raman-spectroscopy or near infrared-spectroscopy (25, 26) introduce additional complexity with regard to data handling, model calibration and analysis (27). A typical approach to overcome this issue is the implementation of enzymatic assays for at-line determination of metabolites and substrates. However, additional equipment like a second LHS or a plate reader is necessary for sample processing and measurements (28), while extensive hands-on time is required for the implementation and validation of the methods. In contrast to that, high-throughput analyzers (HT-analyzer) offer a variety of ready-made assay kits including standard calibration and validation procedures. Due to their large dimensions and bench space in research laboratories being critical, the integration into liquid handling stations becomes difficult (29) especially when cultivation and analytic laboratories are separated. Laboratory automation in these settings remains demanding and usually left unsolved, leading to manual monotonous tasks for the scientist or data gaps have to be accepted. Elegant solutions are needed for linking highthroughput upstream with sample analysis (2), while maintaining the degree of automation. Regarding modular setups for laboratory automation (29) and the connection of different unit operations up to integrated continuous biotechnological processes (30), mobile and collaborative robots are a promising solution. (31) deployed a dexterous mobile robot for automated searching for improved photocatalysts for hydrogen production from water. The robot operated autonomously over eight days with a 1000 times faster workflow compared to manual handling. Although this setup was used in a chemistry research environment, the concepts should also be applicable to biotechnological process development in HT laboratories. The Co-bot technology can be described as a system, which is no longer physically separated but rather integrated with conjunction into the human’s physical work space (32) mainly for improving efficiency, quality and flexibility (33). Other possible tasks are related to ergonomics and safety (32), where the co-bot is mostly responsible for potentially hazardous or monotonous tasks in a time and energy efficient manner (34). Mobile robotic lab assistants allow the integration of different laboratory equipment and different cultivation scales in a common automated platform. Thus, large amounts of necessary process data from different scales can be generated in an automated environment for faster scale-up and reduced timelines (1, 35).

Automated cultivation platforms, mobile robotic technologies and high-throughput analyzers offer potential for flexible and accelerated upstream process development (31). In combination with a centralized data infrastructure model based or data driven process control tools can be easily integrated. In this study, we present the integration of a small scale bioreactor system (≤150 mL) into a LHS in combination with a HT-analyzer. The coupling of the cultivation platform and the HT-analyzer has been realized via a mobile robotic lab assistant, surpassing the challenge of the spatial separation in two separated laboratories. All process data accessible in a centralized database for in-depth process analysis and control. The platform’s key features are (1) automated cultivation control, (2) automated sampling procedure, (3) a mobile robotic workflow for sample transport, (4) at-line quantification of optical density, glucose, acetate and magnesium concentrations, (5) centralized process data and (6) feedback operation loop.

## Material and Methods

### A. BioXplorer Facility

#### A.1. Cultivation System

In this study the BioXplorer 100 (H.E.L group, London, United Kingdom) for eight parallel cultivations in glass STRs with working volumes up to 150 mL was used for this work. The system is equipped with pH sensors (AppliSens, Applikon Biotechnology B.V., Delft, Netherlands), DO sensors (AppliSens, Applikon Biotechnology B.V., Delft, Netherlands), three peristaltic pumps and a mass flow controller for each vessel. The PolyBlock, two columns with four vessels each, holding the vessels was integrated into a Tecan EVO 150 (Tecan Group, Männedorf, Switzerland) equipped with a liquid handling arm with eight steel tips (LiHa), while the control unit of the cultivation system was placed on the left handed side of the LHS. The Pt100 temperature probes for the left reactor column were bend < 30° in accordance with the manufacturer. The temperature profiles of the probes have been tested and were not affected by bending. A cooled MWP carrier was placed on the LHS for storing the sample plates. The process data from the cultivation system e.g. pH, stirring speed, DOT or feed rate was logged in txt-files, as provided by the manufacturer, for each reactor separately. A Python script reads the files and writes the corresponding values to the database. The logging interval was set to 30 s, but a more frequently logging interval can be chosen. Writing set points from the database followed the same principle as reading process data, where a Python script wrote txt-files with corresponding set points for each control loop and reactor individually.

#### A.2. Mobile robotic lab assistant

The mobile robotic lab assistant (Astechproject Ltd., Runcorn, United Kingdom) consisting of a driving platform MIR100, a robotic arm URE5 equipped with a 2-finger gripper and 3D camera was used for automated sample transport. The driving platform is equipped with 3D cameras and laser scanners for navigation and safety. The 2-finger gripper can pick-up MWPs in either portrait or landscape position. Two MWPs can be stored on deck and additional four plates at the back on a shelf of the platform. The laboratory map was taught including device positions for charging station and an additional waypoint. The cultivation system and the high-throughput analyzer were labeled with an april tag and the corresponding positions for the MWP were taught from a right handed position. The corresponding functions for picking and placing a plate were set to linear movements (moveL) in PolyScope 3.2 (Universal robots, Odense, Denmark). The movement velocity of the robot as well as the operating height were restricted due to safety reasons. All necessary functions for setting up and executing a workflow were controlled via a standard in laboratory automation (SiLA2) interface.

### B. High-throughput analyzer

The at-line analysis was conducted by a HT-analyzer (Cedex BioHT, Roche Diagnostics GmbH, Mannheim, Germany), equipped with a rack suitable for 96-MWP and an opening in the front lid (fig. 1C). The following test kits were calibrated and validated with the corresponding controls prior to use and according to the manual: Glucose Bio HT, Acetate V2 Bio HT, Magnesium V2 Bio HT and OD Bio HT (all Roche Diagnostics GmbH, Mannheim, Germany).

**Fig. 1.**
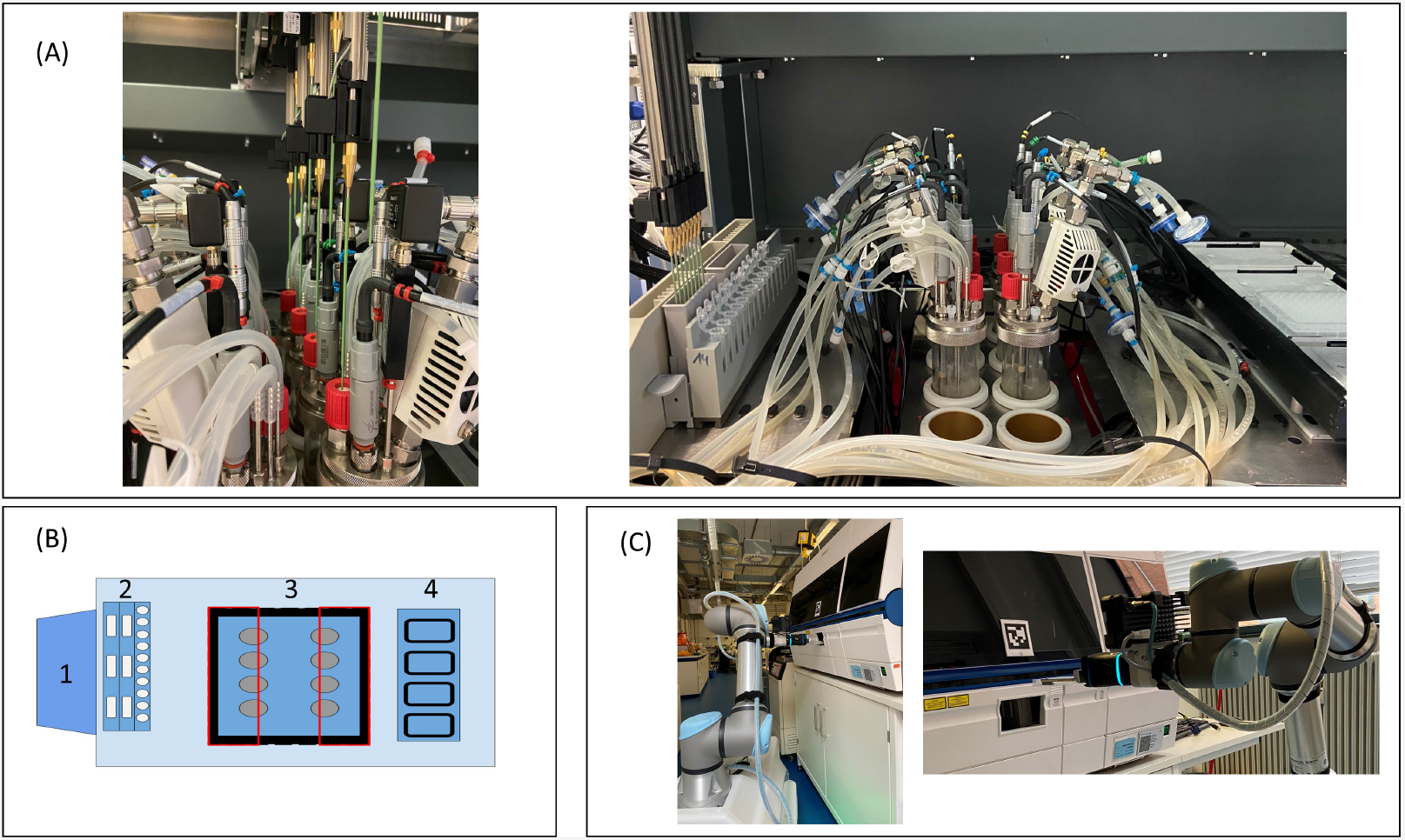
Presentation of the facility. The LHS with the integrated cultivation system **(A)**. Schematic presentation of the integration and LHS deck layout **(B)**. (1) Control station for the cultivation system, (2) wash station, troughs and tube rack, (3) cut-out (black), reactor block (blue), restricted area for the LiHa (red rectangles), (4) cooling rack for 96-MWP. Mobile robotic lab assistant placing the 96-MWP in the HT-analyzer **(C)**. Liquid handling station, LHS; liquid handling arm, LiHa; microwell plate, MWP; HT, high-throughput.

### C. Strain, chemicals, media

All experiments were carried out with *E. coli* BW25113 strain as biological triplicates. All chemicals were obtained from Roth (Carl Roth GmbH, Karlsruhe, Germany), Merck (Merck KgaA, Darmstadt, Germany) or VWR (VWR International, Radnor, PA, USA), if not stated otherwise. TY-medium contained: 16 g L^−1^ bacto tryptone (Becton Dickinson, Franklin Lakes, NJ, USA), 10 g L^−1^ bacto yeast extract (Biospringer,Maisons-Alfort, France) and 5 g L^−1^ NaCl. The main medium consisted of mineral salt medium (MSM), containing: 2 g L^−1^ Na2SO4, 2.468 g L^−1^ (NH4)2SO4, 0.5 g L^−1^ NH4Cl, 14.6 g L^−1^ K2HPO4, 3.6 g L^−1^ NaH2PO4 × 2 H2O, 1 g L^−1^ (NH4)2-H-citrate and 1 mL antifoam (Antifoam 204, Sigma-Aldrich, St. Louis, MO, USA). The medium was supplemented with with 2 mL L^−1^ trace elements solution and (1 M) MgSO4 solution. The trace element solution comprised: 0.5 g L^−1^ CaCl2 × 2 H2O, 0.18 g L^−1^ ZnSO4 × 7 H2O, 0.1 g L^−1^ MnSO4 × H2O, 20.1 g L^−1^ Na-EDTA, 16.7 g L^−1^ FeCl3 × 6 H2O, 0.16 g L^−1^ CuSO4 × 5 H2O, 0.18 g L^−1^ CoCl2 × 6 H2O, 0.087 Na2SeO3 g L^−1^ (Alfa Aesar, Haverhill, MA, USA), 0.12 Na2MoO4 × 2 H2O, 0.725 Ni(NO3)2 × 6 H2O. The feed solution contained 600 g L^−1^ glucose dissolved in MSM and supplemented with 2 mL L^−1^ trace elements solution. For the automated magnesium addition 500 mM MgSO4 solution was used.

### D. Cultivation

#### D.1. Precultivation

15 mL of TY-medium was directly inoculated with 100 μL of cryoculture and cultured in a 125 mL Ultra Yield flask (Thomson Instrument Company, Oceanside, CA, USA) sealed with an AirOtop enhanced flask seal (Thomson Instrument Company, Oceanside, CA, USA) for 7 h at 37 °C and 220 rpm in an orbital shaker (25 mm amplitude, Adolf Kühner AG, Birsfelden, Switzerland). The second preculture was set to an OD_600_ of 0.25 and cultured in 50 mL EnPresso B medium (Enpresso GmbH, Berlin, Germany) with 9 U L^−^1 Reagent A in 500 mL Ultra Yield flask sealed with an AirOtop enhanced flask seal under the same conditions. This allows for continuous glucose release over time and prevents overfeeding. After 12 h appropriate volumes of the preculture were used to inoculate the main culture to an OD_600_ of 0.5.

#### D.2. Parallel STR cultivation

Cultures were run in sixfold parallel glass stirred tank reactors (STR) each equipped with one Rushton type impeller at 37 °C and pH was controlled at 7.0 with 10 % (v v^−1^) NH_3(*aq*)_ via the WinISO control software (H.E.L group, London, United Kingdom) The main cultures were started as 90 mL batch cultures with an initial glucose concentration of 10 g L^−1^. After the batch phase ended, a fed-batch was started with an exponential feeding rate μ_*set*_ = 0.2 or μ _*set*_ = 0.15 for 6 h or 13 h, respectively. Afterwards, the feed was set to constant until the end of the cultivation. The system was aerated with compressed air from 0.5 vvm during the batch phase up to 2 vvm during the fed-batch phase. Additionally, stirring was increased from 1000 rpm during the batch phase up to 1500 rpm during the fed-batch phase, following preset values in WinISO. Manual adjustment to stirring or aeration were conducted, if necessary.

### E. Sampling and analytics

Automated sampling was performed and scheduled by the liquid handling station using a Freedom EVOware (Tecan Group, Männedorf, Switzerland). For the sampling procedure an adapted liquid class with an aspiration speed of 50 μL s^−1^, dispense speed of 150 μL s^−1^ and liquid density of 1 mg mL^−1^ was used. Upon each sampling event the tips were washed and sterilized with ethanol. The left and the right column of the PolyBlock were sampled sub-sequentially and each reactor was sampled through its septum. The sampling triggered the creation of an unique sampling ID for each bioreactor in the database, including the corresponding sampling method (at-line or off-line), timestamp and sampling volume. For off-line measurements 1000 μL per reactor were taken and 2 × 500 μL were dispensed in 1.5 mL reaction tubes. For at-line sampling 600 μL per reactor were taken. The at-line sample volume was dispensed on a 96-MWP in 3 columns à 200 μL. Samples for at-line analysis were inactivated directly with NaOH and stored on 96-well plates at 4°C (28) on the deck of the LHS. The remaining sample volumes after at-line analysis were frozen at -20 °C. Glucose, acetate, magnesium and OD_583_ were analyzed by the high-throughput analyzer. Samples for off-line cell dry weight measurements were collected in 1.5 mL pre-weighed Eppendorf tubes. The tubes were centrifuged at 4 °C, 10000 rpm (Hitachi Koki Co. Ltd., Tokyo, Japan) for 10 min. The supernatant was discarded, the pellet was dried at 85 °C for 24 h, acclimated in a desiccator and weighed. The CDW was calculated from the mass difference and the collected values were written to the database. Measurements were conducted as duplicates.

## Results

The presented study illustrates the integration of a small scale robotic cultivation system and a HT-analyzer via implementing a mobile robotic lab assistant in the existing facility. All process data was stored in a centralized database. As a case study *E. coli* BW25113 high cell density cultivations were performed. Necessary magnesium addition steps to MSM, during HCD cultivations, were automated via a feedback operation loop.

### F. Device integration and data handling

For the physical device integration, adaptations regarding the stainless steel deck of the LHS were necessary. The deck has been removed and in order to fit the reactor block into the LHS, a customized cut-out in the deck has been realized. Therefore, the PolyBlock is located below the LHS deck level, guaranteeing the necessary travel height for the steel tips over the reactors (fig. 1A). As demonstrated in figure 1B the area above the cut-out was divided into two inaccessible areas, highlighted as two rectangles, allowing the LiHa only to move in a defined vertical line to reach the septum ports. Thus, preventing collisions of the LiHa with one of the probes, the condensers or any other tubing on the head plate. These modifications enable automated sampling through the septum port of each reactor with a minimum culture volume of ca. 80 mL via the LiHa. As shown in figure 1C the HT-analyzers front lid was equipped with a customized opening by the manufacturer. The opening grants the robot arm access to a 96-MWP rack without manually opening and closing the lid.

The data integration followed the concepts presented by (6) ensuring that all experimental data for a run is saved in a centralized and accessible manner. A schematic representation of the device and data integration is depicted in figure 2. The reading and writing procedures for process data, set points and control actions were automated via Python scripts (fig. 2). The set points of the cultivation system can be either controlled via the control software WinISO or via the centralized database. No propriety data format is needed. A basic implementation for controlling temperature, pH, air flow rate, stirring and feed pump rates was developed and can be easily connected to other interfaces. A Python based GUI was developed, in order to facilitate the control of the data logging and set point writing. By that the cultivation system can be controlled flexibly via set points from the database or via the manufacturer’s control software (WinISO). The automated sampling procedure conducted by the LHS was scheduled via the liquid handling script (Freedom EVOware). During each sampling event an entry in the database is created featuring the sampling method (at-line or off-line) the sample volume, the timestamp and an unique sample ID. After the transfer of the samples from the cultivation system to the HT-analyzer, those sample IDs were used as label names for the samples. Since the sampling procedure and the at-line measurements have different timestamps, the measurements were matched to the actual timestamp of the sampling procedure. According to that, a Python script pulls the corresponding timestamps for each sample ID from the database and matches the measured values for glucose, acetate, OD_583_ and magnesium. Thereby, all on-line and at-line process data is accessible in the database and can be deployed by other applications e.g. feedback operation, model based control or machine learning algorithms. As an example for such a feedback operation loop, magnesium addition (see H.1) was automated. The feedback operation loop was implemented as a sub-method in a modular way. Hence, the setup can be easily adjusted for different control or trigger based applications e.g. automated induction. The control procedure was initiated by the LHS script and the magnesium controller calculated the corresponding volumes to be added, based on the set points and recent process data in the database. The necessary pipetting steps for each reactor were written to a worklist (gwl-list). The gwl-list contained the pipetting instructions, which was loaded and subsequently executed by the LiHa.

**Fig. 2.**
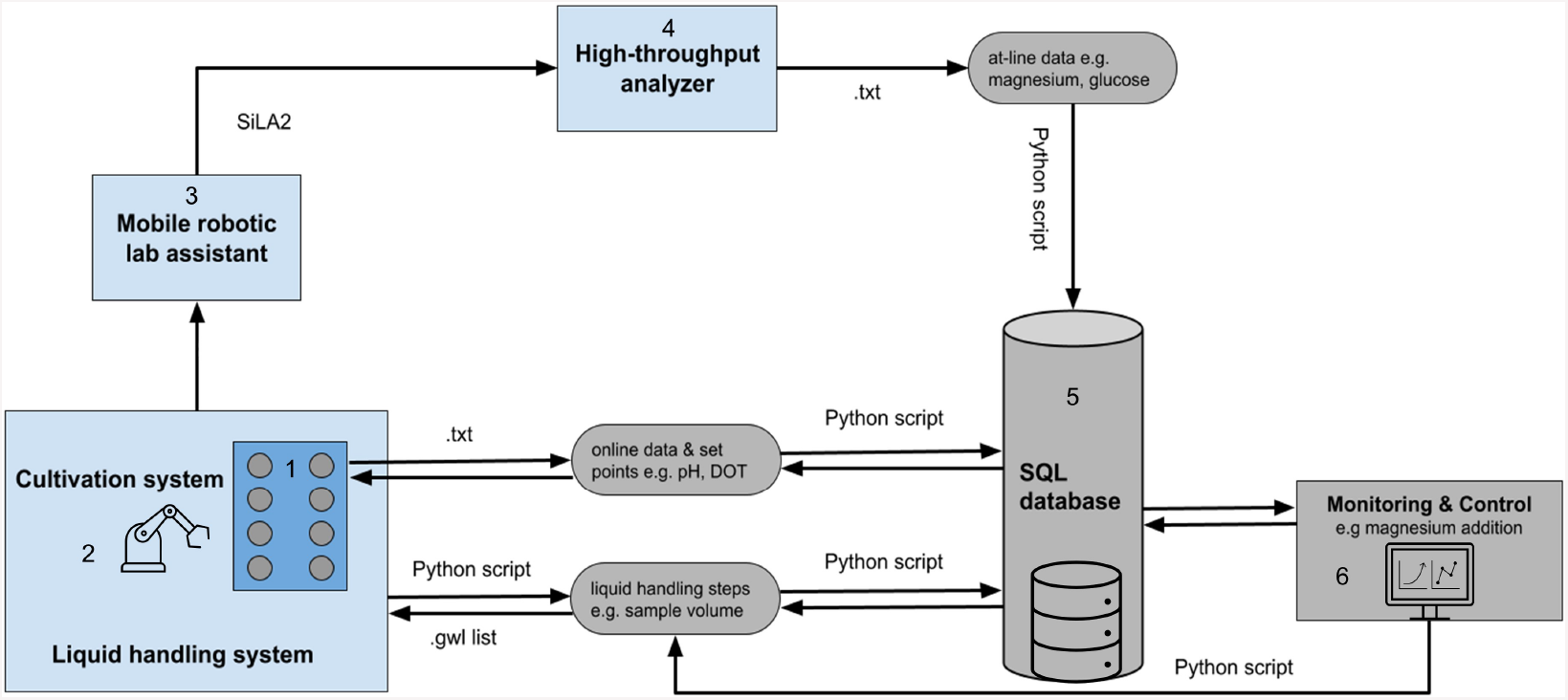
Schematic representation of the device (light blue) and data integration (light grey) in the facility. The facility is composed of a (1) Cultivation System, (2) Liquid handling station, (3) Mobile robotic lab assistant, (4) High-throughput analyzer, (5) SQL-database and a (6) Monitoring and Control system. All process data is stored in the central database. The communication, data transfer and control relies on different file formats e.g. txt- and gwl-files. Necessary reading and control dependent writing of these files are automatically performed via Python scripts as well as database queries. The mobile robotic lab assistant is controlled via a SiLA2 server and client architecture. Structured Query Language, SQL; Standard in Laboratory Automation, SiLA.

### G. Mobile robotic lab assistant

The HT-analyzer and the cultivation platform are spatially separated in two different rooms, the robotic cultivation laboratory and the analysis laboratory. The spatial separation of devices to be used inevitably leads to a decrease in the degree of automation. To overcome this issue and maintain a certain degree of automation a mobile robotic lab assistant was integrated in the facility. The map, including overhand and device positions (6B), was taught beforehand as described in section A.2. Different workflows were tested for transferring the 96-MWP containing the samples from the cultivation system to the HT-analyzer. The main obstacles for an efficient workflow were identified and the following points were addressed: (1) restrictions in laboratory space, (2) restrictions of the robotic arm, (3) restrictions of integrated laboratory instruments.

The transport of the 96-MWP required the mobile robotic lab assistant to move from the cultivation laboratory to the analysis laboratory. Research laboratories suffer from available space and empty bench space can be a limiting restriction for the location of equipment. The HT-analyzer is located in the middle between two work-benches. To access the HT-analyzer the robot had to navigate through a narrow corridor between these work-benches. In combination with laboratory staff being around, the direct routing from the cultivation system to the HT-analyzer was prone to fail. Instead of going directly from the cultivation system to the HT-analyzer an additional stop, referred to as way-point, was incorporated in the workflow. The way-point area was kept free from other objects. Starting the routing through the narrow corridor from the way-point prevented routing errors of the driving platform.

The robotic arm is equipped with a 2-finger gripper and a mounted 3D camera (6A). The 360° radius of the robotic arm allows to teach and approach devices from a left-handed or right-handed position. Due to the configuration of the robotic arm and the position of the joints, a left-handed or right-handed movement leads to different angles and joint positions of the robotic arm (6C,D). Depending on the task that has to be performed left-handed or right-handed movements were restricted. Especially, left-handed arm movements have been difficult regarding restrictions of the laboratory devices. In order to circumvent collisions, picking and placing of 96-MWPs was carried out from a right-handed position. The HT-analyzer is equipped with an opening at the front lid (fig. 1C). The opening allows the robotic arm to access the HT-analyzer and place or pick the 96-MWP on the sample rack. Due to the narrow dimensions of the opening, it was necessary to avoid any side-ward movements of the robotic arm, while placing or picking a 96-MWP. To guarantee straight robotic arm movements, the corresponding methods for placing and picking the 96-MWP were set to linear (see A.2). The approach distance, that the robot arm will move to before picking or placing the 96-MWP, was kept at the maximum of 300 mm. Hence, already from that distance on the movements were kept linear. The final workflow for the sample transfer is represented in figure 3 and consisted of the following steps: (1) move 96-MWP from LHS to mobile robotic lab assistant, (2) place new 96-MWP from mobile robotic lab assistant’s back-shelf to LHS, (3) go to way-point, (4) go to HT-analyzer, (5) place 96-MWP on rack in HT-analyzer, (6) remove 96-MWP from HT-analyzer, (7) go to way-point and (8) stay idle.

**Fig. 3.**
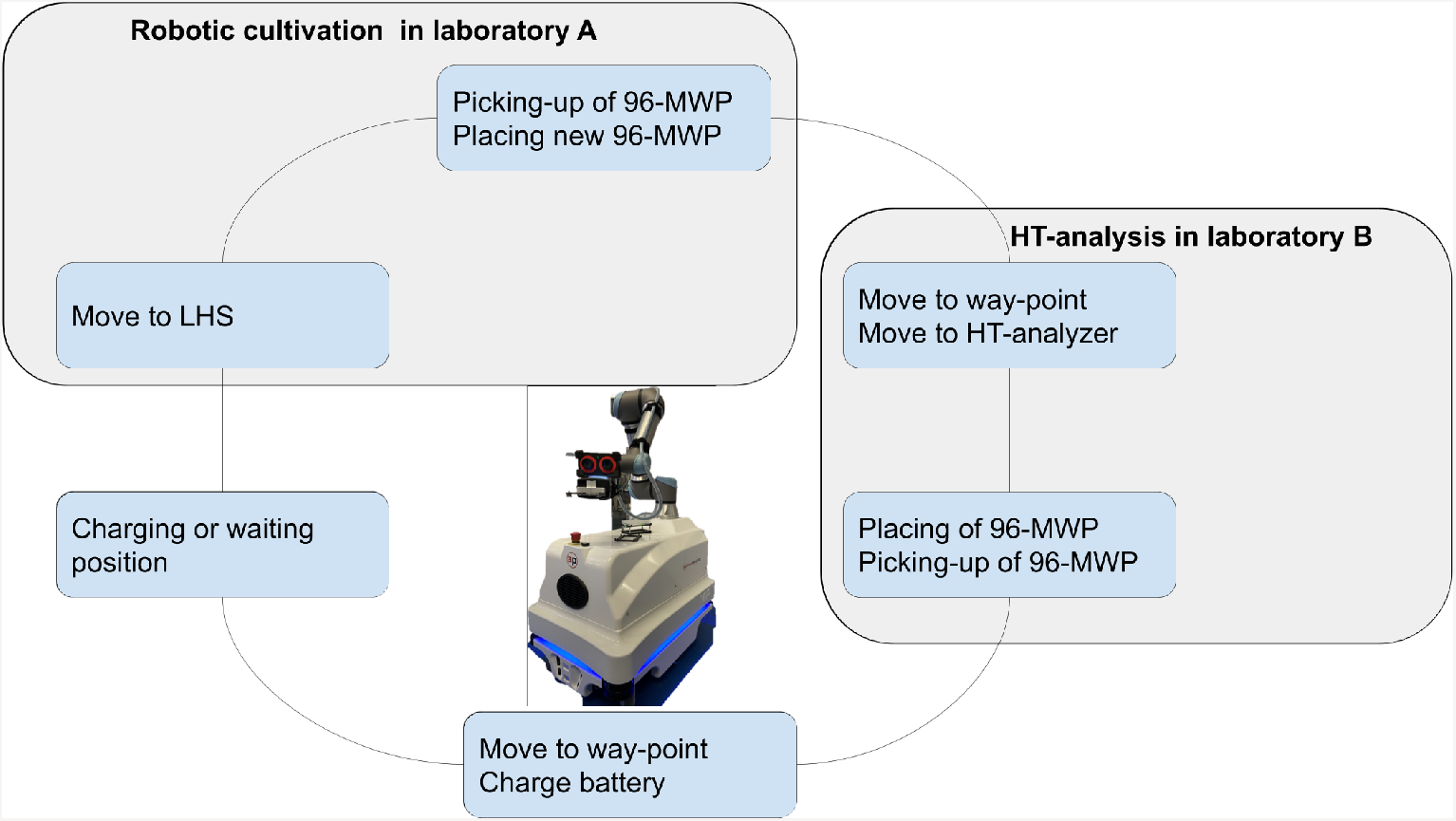
Illustration of the mobile robotic workflow. The mobile robotic lab assistant connects the cultivation platform in laboratory A with the high-throughput analyzer in laboratory B. The established workflow for the transportation of 96-microwell plates consisting of the following steps:(1) move 96-MWP plate from the LHS to mobile robotic lab assistant, (2) place new 96-microwell plate from mobile robotic lab assistant back-shelf to LHS (3) go to way-point (4) go to HT-analyzer (5) place 96-MWP on rack in HT-analyzer (6) remove 96-MWP from HT-analyzer (7) go to way-point (8) charge battery. Microwell plate, MWP; high-throughput, HT; liquid handling station, LHS.

### H. Proof of concept high cell density cultivation

Industrial cultivations are mostly performed in fed-batch mode aiming for high cell densities, where the limited substrate is continuously fed with a certain rate. Thus negative effects of overflow metabolism can be avoided and high cell densities can be reached. The developed system was evaluated based on its capability of performing an industrial relevant HCD cultivation. As a proof of concept *E. coli* BW25113 cultivations were designed with the intention of reaching high cell densities. Hence, two feeding profiles were compared as biological triplicates. Both profiles followed an industrially relevant feeding regime starting with a batch phase followed by an exponential feed and constant feed and were designed to reach approximately 50 g L^−1^ CDW within 30 h of cultivation.

For adequate process monitoring and control, online as well as at-line data was logged in the central database as described above (see F). The on-line process data such as DOT, pH, stirrer speed, air flow and cumulative volumes for base and feed addition are shown in the supplementary material (7). The measurements for OD, glucose, acetate, CDW as well as the feeding profile are shown in figure 4. Throughout the cultivation OD (fig. 4A,B), glucose (fig. 4C,D) and acetate (fig 4E,F) were analyzed at-line in the HT-analyzer. The biological replicates with an exponential feeding rate of μ_*set*_=0.2 h^−1^ are referred to as reactor 1-3 (R1-R3) and biological triplicates with an with an exponential feeding rate of μ_*set*_=0.2 h^−1^ are referred to as reactor 4-6 (R4-R6). The reactors were inoculated to an OD_600_ of 0.5 in mineral salt medium with an initial glucose concentration of 10 g L^−1^. Throughout batch phase of the cultivation OD was measured (fig. 4A,B), allowing to monitor the growth at-line. Based on that a maximum specific growth rate of 0.57 +- 0.024 h^−1^ was determined (R1 excluded). During the batch phase the glucose depleted for each of the triplicates (fig. 4C,D). R5 showed a higher initial glucose concentration of 13.37 g L^−1^ compared to the others (fig. 4D). While glucose depleted a typical increase in the acetate concentration, due to overflow metabolism, can be seen. The acetate concentration increased for R2-R6 up to values between 0.5 and 0.76 g L^−1^ (fig. 4E,F). The following acetate re-consumption lead to a decrease in the acetate concentration and is also shown in the pH peak. For R1 acetate increased up to 2.38 g L^−1^ (fig. 4E) due to a stirrer malfunction and insufficient mixing. After the glucose was consumed and the corresponding peak in the DOT signal appeared between 5.6 and 6.3 h of cultivation, the fed-batch was started. The exponential feeding rate was set to μ_*set*_=0.2 h^−1^ for 6 h and μset_*set*_=0.15 h^−1^ for 13 h for R1, R2, R3 (fig. 4G) and R4, R5, R6 (fig. 4H), respectively. Glucose limitation was maintained throughout the exponential feeding phase and overfeeding was avoided. However, after 22 h during the constant feeding phase, glucose and acetate concentrations increased for all reactors. R1 showed a maximum accumulation of glucose and acetate with 1.22 g L^−1^ (fig. 4C) and 0.371 g L^−1^ (fig. 4E), respectively. The constant feeding rate for R6 was reduced after ca. 27.5 h (fig. 4H) in order to avoid oxygen limitation. In addition to atline measurements a sampling procedure for off-line CDW measurements was automatically started during the feeding phase. Figure 4I,J shows the off-line CDW measurements for the biological triplicates. The biomass steadily increased during the feeding phase for R1-R3 (fig. 4I) as well as for R4-R6 (fig. 4J). A gap in the off-line measurements between 17 h and 26 h of cultivation was due to an intended sampling break during the night. After 26 h of cultivation R4 showed a lower biomass of 36.2 g L^−1^ ±1.8 g L^−1^ compared to R5 and R6. At the end of the cultivation R4 stayed below the final 50 g L^−1^ with final biomass of 43.9 g L^−1^ ± 5.8 g L^−1^. Besides R4 (fig. 4I), the desired CDW of 50 g L^−1^ was reached for both feeding profiles. It can be seen from the shown data, that the variance of the biological triplicates is comparatively low and the error of the at-line measurements is negligible.

**Fig. 4.**
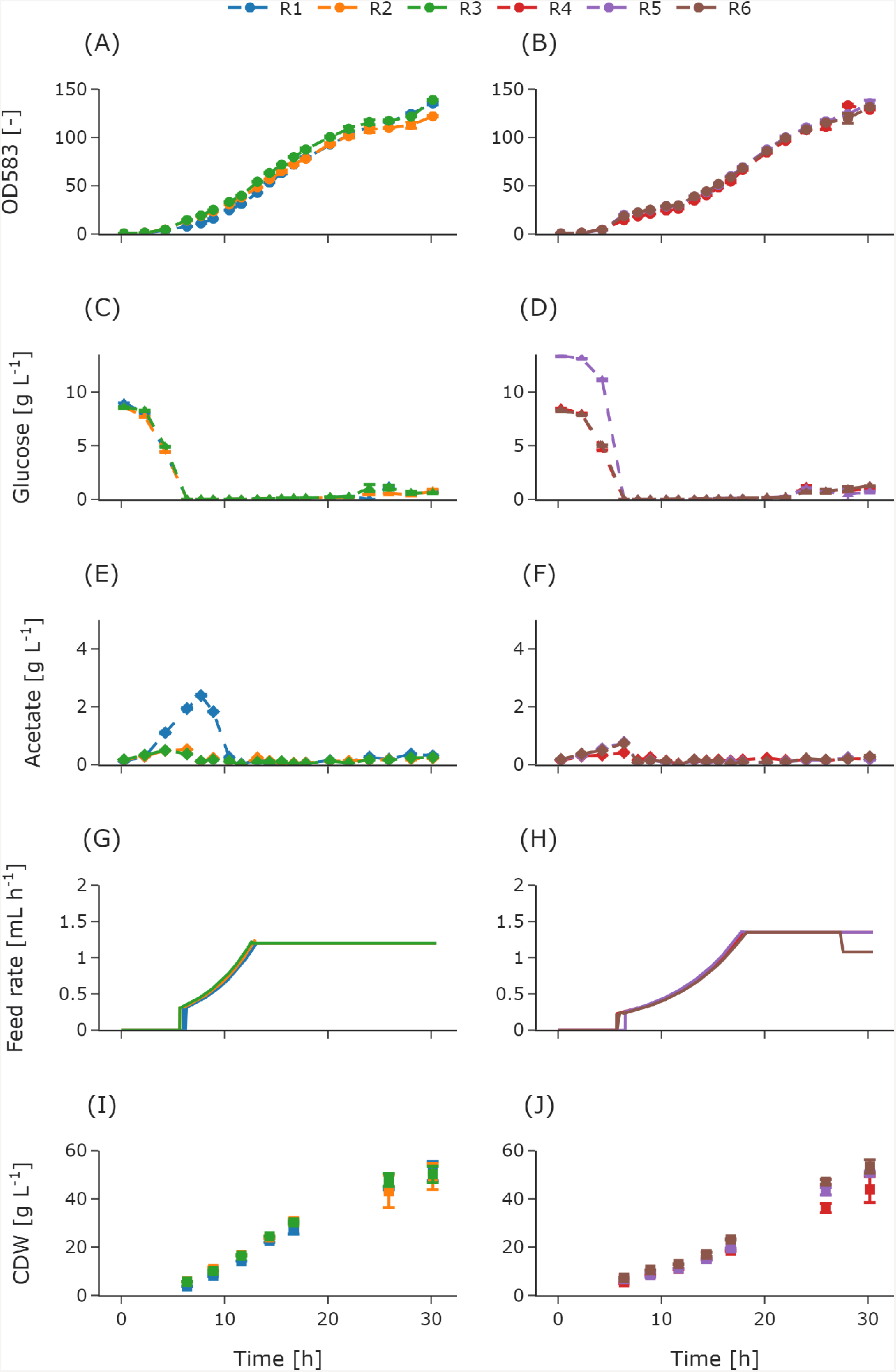
Measured at-line and off-line values of *E. coli* BW25113 fed-batch cultivations with two different feeding profiles as biological triplicates. OD_583_ for R1-R3 **(A)** and OD_583_ for R4-R6 **(B)**, glucose for R1-R3 **(C)** and glucose for R4-R6 **(D)**, acetate for R1-R3 **(E)** and acetate for R4-R6 **(F)**, feed rate for R1-R3 **(G)** and feed rate for R4-R6 **(H)** CDW for R1-R3 **(I)** and CDW for R4-R6 **(J)**. For R1-R3 the exponential feed was set to μ_*set*_=0.2 h^−1^ for 6 h followed by a constant feed. For R4-R6 the exponential feed was set to μ_*set*_=0.15 h^−1^ for 13 h followed by a constant feed. Error bars derived from replicates (n=2). Reactor, R; optical density, OD, cell dry weight, CDW.

#### H.1. Feedback operation

Automation of parallel experiments requires additional control loops such as trigger based events or feedback operation loops. In this study, magnesium addition was automated to demonstrate the capability of the system to perform feedback operation loops. The feedback operation loop was implemented, based on the magnesium measurements from the HT-analyzer. The implementation is described in section F. Mineral salt medium requires the addition of magnesium to the batch medium. During the fedbatch phase the feed solution can be supplemented (13) or manual addition of magnesium solution is necessary. The magnesium set point for the control loop was 2 mM based on the concentration in the batch medium and in accordance with the lower limit of the used test kit. Figure 5 depicts the measured magnesium concentrations (fig. 5A) and the added volumes (fig. 5B) during the cultivation for all reactors. The initial measured values were below 2 mM. Hence, the control loop initiated the first magnesium addition cycle via the LHS already during the batch phase. As demonstrated in figure 5B the corresponding volumes were tracked for each reactor individually. Throughout the batch phase magnesium was added automatically in every control cycle to all reactors. For the following exponential feeding phase, magnesium was also added in every control in order to reach the set point of 2 mM. When the feed was switched to constant after 6 h of exponential feed for R1, R2 and R3, the magnesium concentration reached the set point of 2 mM (fig. 5A). Therefore, no further magnesium was added to these reactors in the upcoming control cycles and the cumulative volume stayed the same 5B. The same behaviour was observed 7 h later, when the constant feed for R4, R5 and R6 started. Towards the end of the cultivation the magnesium concentration in the medium increased for all reactors (fig. 5A). Consequently, the feedback operation loop stayed inactive. The total amount of magnesium solution added varied between a minimum of 1.55 mL for R1 and maximum of 2.1 mL for R2 (fig. 5B).

**Fig. 5.**
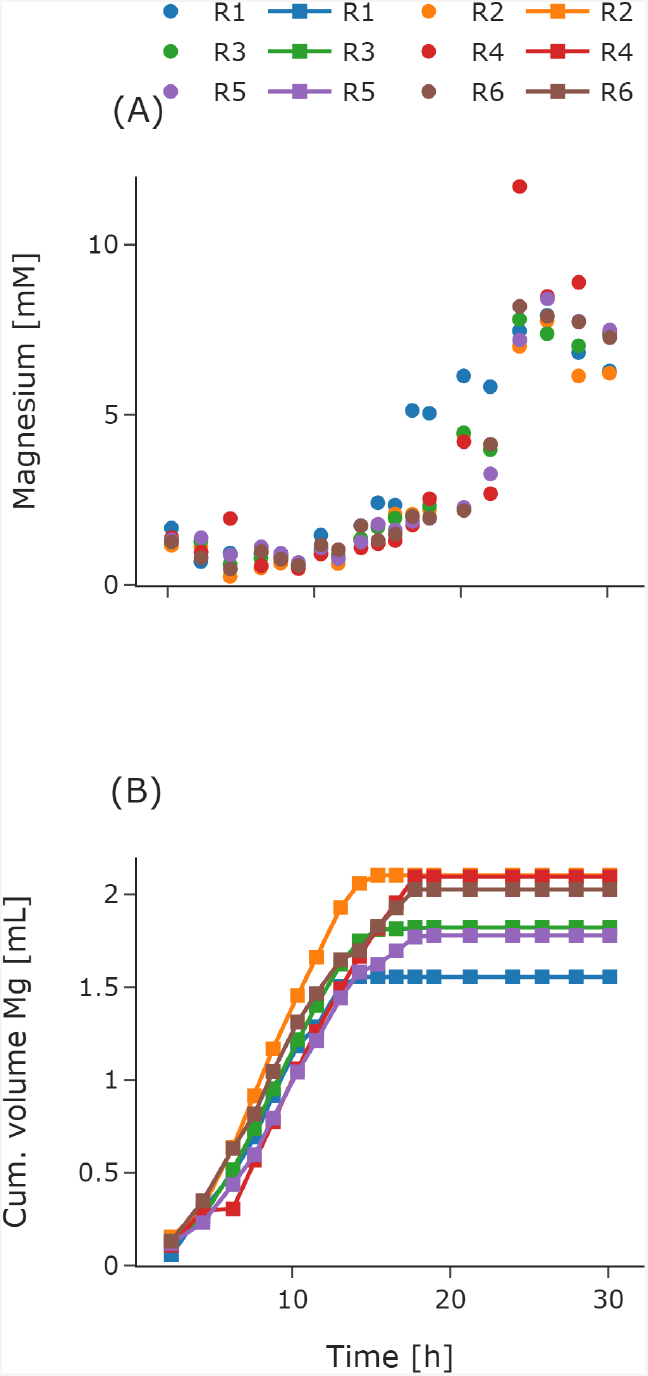
Automated feedback operation for magnesium addition. At-line measured magnesium concentrations throughout the cultivation for R1 - R6 **(A)**. Cumulative volume of the added magnesium solution for R1 - R6 **(B)**. The volume for each control cycle was calculated according to section H.1 with a set point of 2 mM. For R1-R3 the exponential feed was set to μ_*set*_=0.2 h^−1^ for 6 h followed by a constant feed. For R4-R6 the exponential feed was set to μ_*set*_=0.15 h^−1^ for 13 h followed by a constant feed. Reactor, R.

## Discussion

Automated cultivation platforms require the integration of different devices. The coupling of devices is restricted by large footprints and available laboratory space (29), limiting automated cultivation workflows. In this study we addressed this challenge and integrated a robotic cultivation system and a spatially separated HT-analyzer. The mobile robotic lab assistant coupled the devices via automated sample transfer. All process data was stored in a centralized database and magnesium addition was automated via the feedback operation loop.

The integration of cultivation systems into LHS is a well-established approach for automated operation and sampling procedures and has been addressed before (4, 6, 7). Nevertheless, it lays the foundation for automated cultivation workflows and further automation steps. The presented robotic cultivation system allowed for automated liquid handling procedures. Additional sampling ports or increased sampling volumes due to dead volumes were avoided. To gain process relevant insights, at-line analytics were conducted on a HT-analyzer. The HT-analyzer offers a variety of pre-made assay kits with standard procedures for calibration and validation. No extra hands-on time for implementation and validation of the enzymatic assays were needed. Hence, the atline capabilities can be easily extended with reduced effort. The on-line, at-line and off-line data was stored in a centralized database. The data handling was successfully automated via Python and allowed for flexible control of the cultivation system. The demand for transferring automation tools from microbioreactor systems to laboratory scale cultivation systems has recently been addressed (18). In this study, the data integration as well as the modular design of the feedback operation loop followed the principles of our existing HT bio-process development facility (6). In addition to transferring automation tools, the connection to the centralized database sets the foundation for an integrated facility with different cultivation scales.

In comparison to existing robotic facilities (4, 6, 7, 18), we addressed the physical integration for at-line analytics in a substantially different way. A mobile robotic lab assistant controlled via a SiLA2 interface connects the cultivation platform with a HT-analyzer for automated sample transport with subsequent at-line analysis. Restrictions regarding the robotic arm, the integrated laboratory devices and the available space in the laboratories were identified. The established workflow handled these restrictions and circumvented changes in the laboratory layout. As emphasized by (29) flexible automation tools are an important factor for research laboratories with varying experimental protocols. A modular design approach allows for flexible automation according to the current protocol. Thus avoiding the purchase of redundant features and allowing possibilities for adaptations. The demonstrated setup has specific advantages with regard to flexibility and modularization. The mobile robotic lab assistant successfully enabled the coupling of the cultivation platform to a HT-analyzer in a different laboratory. Thus, spatial separation of laboratory devices is no restriction in this setup, neither are the analytics restricted to one dedicated device. The robotic workflow can be flexibly extended in order to integrate further equipment up to connecting different unit operations from upstream to downstream processing.

As highlighted by (9) industrial scale processes are usually performed as HCD fed-batch cultivations. However, high throughput cultivation platforms are either incapable or rarely challenged for high cell densities (16). In this study, HCD cultivations were performed for accessing the feasibility of the developed system. The results demonstrate the successful execution of parallel high cell density cultivations. A final CDW of ca. 50 g L^−1^ was reached with two different feeding profiles. R4 reached only a final CDW of 43.9 g L^−1^ ± 5.8 g L^−1^. This might due to a handling error during the off-line procedure as indicated by the large error bar. The atline OD measurements of the biological triplicates (fig. 4B) rather suggest a similar final biomass. The gathered at-line data showed low standard deviation emphasizing a robust operation of the system and allowed for good monitoring of the process. During the exponential feeding phase, no glucose accumulation was observed and overfeeding was avoided for both feeding rates respectively. The slight accumulation of glucose towards the end of the cultivation is likely due to a decreasing glucose uptake rate as typically seen with decreasing specific growth rates towards the end of cultivations (36). The acetate concentration remained below 0.4 g L^−1^ throughout the feeding phase. This is comparable to other HCD cultivations where specifically the production of overflow metabolites was supposed to be avoided (37, 38). Hence, the proposed setup is suitable for operating, monitoring and controlling HCD cultivations in an appropriate manner.

The established robotic workflow enabled the acquisition of at-line measurements throughout the cultivation avoiding gaps in the data due to manual handling steps as exemplified by the off-line CDW measurements (see. H). The consistently gathered at-line data was used for further automation steps. A feedback operation loop for automated magnesium addition was successfully implemented and allowed for individual magnesium addition to each reactor. The feedback operation demonstrated in this study serves as a blueprint and is interchangeable for any other at-line measurement and subsequent control procedure. As a central data infrastructure is given via the database, model-based tools e.g adaptive feeding strategies (21) or model predictive control (39) can be transferred and deployed. Thus, fundamental prerequisites towards a data driven platform for accelerated bioprocess development are set.

Although the data infrastructure with a centralized database as well as the mobile robotic lab assistant allow for automated workflows, obstacles hindering a full autonomous operation remained unaddressed. The HT-analyzer interface for automatically starting and stopping measurements is such an obstacle. For rapid and flexible integration of laboratory devices, standard protocols and unified data formats for plug and play operation are required (40). Such concepts provide a solution towards a fully automated workflow. Nevertheless, the conceptual design of the platform aligns well with described demands for accelerated (9) and consistent bioprocess development (1)like modularization, integration of robotic technologies, fed-batch operation or advanced control algorithms such as modelling tools.

In conclusion, the platform closes a bottleneck between HT automated robotic screening facilities and individually controllable laboratory scale reactors, transferring the automation tools to small scale reactors with higher process flexibility, individual control and continuous feeding. Through the operation under industrial relevant process conditions, risk for failure during development and scale-up can be reduced. The proposed setup including the implementation of a mobile robotic lab assistant into the overall workflow is well applicable to other laboratories, where spatial separation of devices or unit operations from upstream to downstream restricts automation. This study provides the basis for a fully automated facility that allows the integrated development of process steps of different scale and sequence. Thus, by linking the individual process steps, the acceleration of the entire bioprocess development pipeline can be achieved.

## ACKNOWLEDGEMENTS

We would like to thank Simon Seidel and Annina Kemmer for their database connection modules. We would like to thank Niels Krausch and Fabian Schröder-Kleeberg for the introduction into the facility. We thank Roche CustomBiotech (Mannheim, Germany) for the supply of the Cedex Bio HT Analyzer. We acknowledge support by the Open Access Publication Funds of TU Berlin.

## COMPETING FINANCIAL INTERESTS

Sebastian Groß is employed by wega Informatik GmbH. M.-Nicolas Cruz-Bournazou is employed by DataHow AG. The remaining authors declare that the research was conducted in the absence of any commercial or financial relationships that could be construed as a potential conflict of interest.

## AUTHOR CONTRIBUTIONS

Conceptualization: LK, SG, MTS, MNCB; methodology: LK, SW; software: LK; validation: LK, SW, SR; formal analysis: LK; investigation: LK, SG, MTS; resources:LK, SW; data curation: LK; writing|original draft preparation: LK; writing|review and editing: SW, MTS, SR, MNCB, PN, SG; visualization: LK; supervision: MTS, SR, SG; project administration: MNCB; funding acquisition: PN. All authors have read and agreed to the published version of the manuscript.

## FUNDING

This work was supported by the German Federal Ministry of Education and Research through the Program “International Future Labs for Artificial Intelligence” (Grant number 01DD20002A)

## DATA AVAILABILITY STATEMENT

The Python code for controlling the cultivation system remotely by other applications can be found in [HEL_database_connector] [https://git.tu-berlin.de/bvt-htbd/facility/hel_database_connector]. The Python code for the feedback operation loop can be found in [BioXplorer_Tecan] [https://git.tu-berlin.de/bvt-htbd/facility/bioxplorer_tecan]. Access to the repositories can be granted upon request.

## Figures and Supplemental Data

**Fig. 6.**
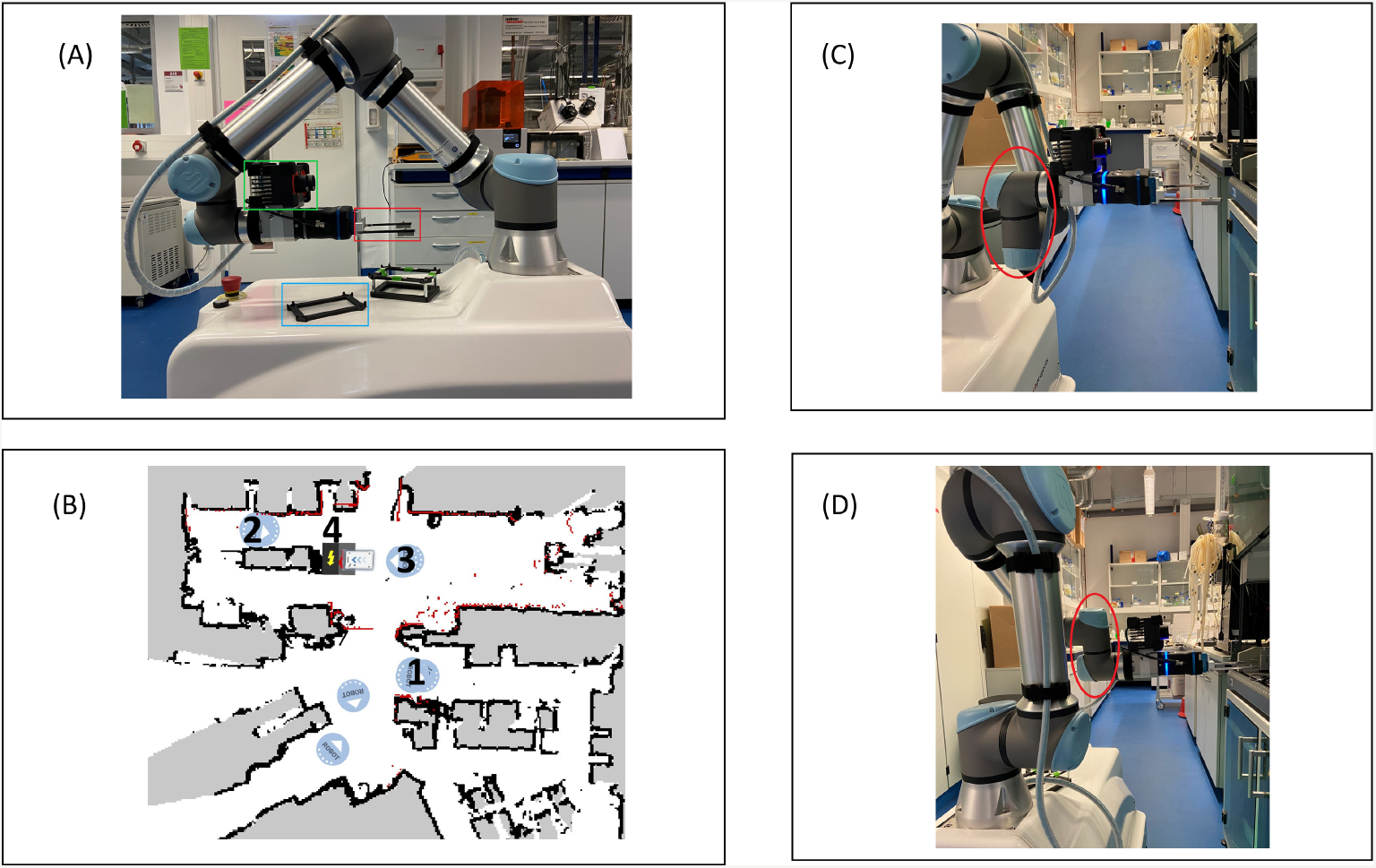
Supplemental:Mobile robotic lab assistant. Mobile robotic lab assistant with 3D camera mounted on the robotic arm (green rectangular), 2-finger gripper (red rectangular) and on deck plate storage (blue rectangular) **(A)**. The taught map with device positions for the (1) cultivation platform, (2) HT-analyzer, (3) way-point, (4) charger **(B)**. Robotic arm movement towards a right-handed position **(C)** and a left-handed position **(D)** with different joint configurations (red circles).

**Fig. 7.**
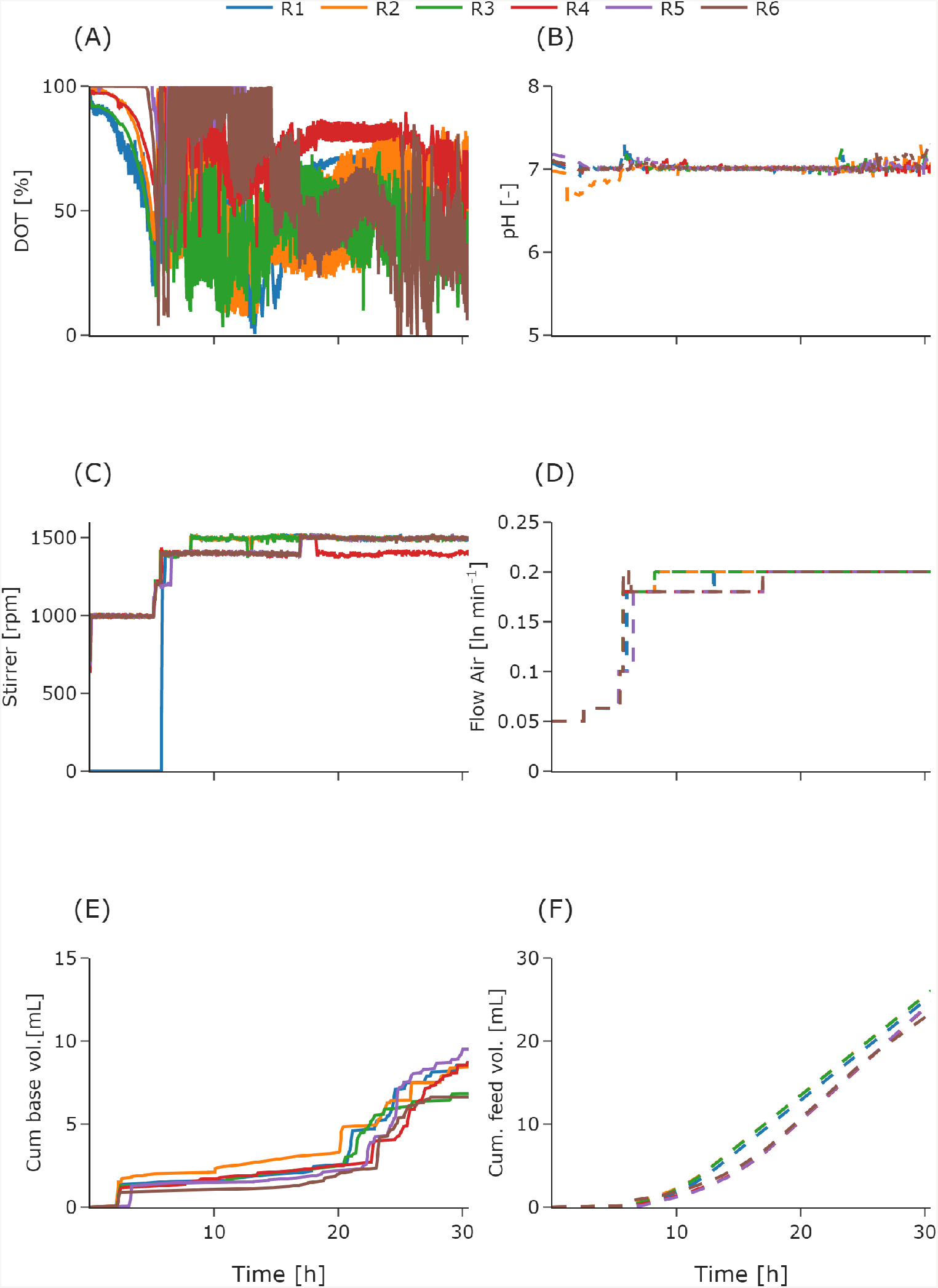
Supplemental:On-line process data of *E. coli* BW25133 fed-batch cultivation’s with two different feeding profiles as biological triplicates. For R1-R3 the exponential feed was set to μ_*set*_=0.2 h^−1^ for 6 h followed by a constant feed. For R4-R6 the exponential feed was set to μ_*set*_=0.15 h^−1^ for 13 h followed by a constant feed. DOT **(A)** and pH **(B)**, stirrer speed**(C)** and flow air **(D)**, cumulative base volume **(E)** and cumulative glucose feed volume **(F)** were measured online.

## Notes

https://git.tu-berlin.de/bvt-htbd/facility/bioxplorer_tecan

https://git.tu-berlin.de/bvt-htbd/facility/hel_database_connector

### Competing Interest Statement

Sebastian Gross is employed by wega Informatik GmbH. M.-Nicolas Cruz-Bournazou is employed by DataHow AG. The remaining authors declare that the research was conducted in the absence of any commercial or financial relationships that could be construed as a potential conflict of interest

